# Generating and testing the efficacy of transgenic Cas9 in *Tribolium castaneum*

**DOI:** 10.1101/2021.10.28.466351

**Authors:** Johnathan C. Rylee, Alexandra Nin-Velez, Simpla Mahato, Kennedy J. Helms, Michael J. Wade, Gabriel E. Zentner, Andrew C. Zelhof

## Abstract

CRISPR/Cas9 genome editing has now expanded to many insect species, including *Tribolium castaneum*. However, compared to *Drosophila melanogaster*, the CRISPR toolkit of *T. castaneum* is limited. A particularly apparent gap is the lack of Cas9 transgenic animals, which generally offer higher editing efficiency. We address this by creating and testing transgenic beetles expressing Cas9. We generated two different constructs bearing basal heat shock promoter-driven Cas9, two distinct 3’ UTRs, and one containing Cas9 fused to EGFP by a T2A peptide. For each construct, we were able to generate a line that is homozygous viable, though variable reductions in reproductive success with each construct were noted. Analyses of Cas9 activity in each transgenic line demonstrated that both designs are capable of inducing CRISPR-mediated changes in the genome in the absence of heat induction. Overall, these resources enhance the accessibility of CRISPR/Cas9 genome editing for the *Tribolium* research community and provide a benchmark against which to compare future transgenic Cas9 lines.

## Introduction

*Tribolium castaneum* is a continually evolving genetic model, and *Tribolium* research complements and offers several benefits over the standard and popular insect model of *Drosophila melanogaster*. Not only is *Tribolium* a model for evolutionary and developmental questions of biology ((Adamski *et al*. 2019; Pointer *et al*. 2021)), but also a model for regulating agricultural pests ((Brown *et al*. 2009; Rosner *et al*. 2020)). Moreover, it is a representative of the order Coleoptera, which comprises approximately 25% of animal species as well as 40% of known insect species ((Hunt *et al*. 2007)). While much progress has been made in developing *Tribolium* as a genetic model, it has not yet been widely adopted, possibly due to the paucity of available reagents and lines enabling simple genetic manipulation relative to *Drosophila*.

The genetic tractability of *Tribolium* has been substantially improved using several strategies. *Tribolium* can be genetically manipulated in the lab through insertional mutagenesis using multiple transposable element-based systems (i.e. minos, piggybac, etc.), allowing for the creation of mutant lines and transgene incorporation ((Berghammer *et al*. 1999; Lorenzen *et al*. 2003; Pavlopoulos *et al*. 2004)). RNAi is effective and durable through injection of dsRNA into all life stages, and in the early embryo using parental RNAi ((Brown *et al*. 1999; Bucher *et al*. 2002)). The GAL4/UAS system is effective for controlling transgene expression ((Schinko *et al*. 2010; Rylee *et al*. 2018)). Finally, CRISPR has recently been used by multiple labs to create knock-in and knock-out lines ((Gilles *et al*. 2015; Adrianos *et al*. 2018; Rylee *et al*. 2018; Gilles *et al*. 2019)).

Since the first demonstration of CRISPR/Cas9-mediated genome editing in *Tribolium* ((Gilles *et al*. 2015)), its use has been limited to a handful of reports. In most cases, Cas9 is supplied via injection of a plasmid ((Gilles *et al*. 2015; Rylee *et al*. 2018)) or purified protein ((Adrianos *et al*. 2018)) and leads to mutagenesis or homology-directed repair (HDR). Recently, ReMOT (Receptor-Mediated Ovary Transduction of Cargo) Control ((Chaverra-Rodriguez *et al*. 2018)) was used successfully to target the *Tribolium cardinal* gene ((Shirai and Daimon 2020)). This method of delivery is promising for emerging models because it allows researchers to imprecisely inject materials into adults rather than embryos, thereby saving time and resources needed for training and optimization of embryo injections. However, a major drawback is that ReMOT Control only permits CRISPR/Cas9 mutagenesis; one cannot combine the technique with an exogenous DNA template for the induction of homology directed recombination (HDR). Additionally, while the authors were able to successfully mutate the *cardinal* gene, their reported mutagenesis rates were substantially below those reported in other species ((Shirai and Daimon 2020)).

Whether it’s transposon transposition, PhiC31 mediated insertion or recombination ((Groth *et al*. 2004; Bateman *et al*. 2006)), or CRISPR/Cas9 genome editing, the efficiency and heritability of each process is increased when the key enzyme is supplied endogenously. In addition, limiting enzyme expression to the germline not only increases the chances of germline transmission of the induced edits but reduces potential toxicity to the injected developing embryo. In *Drosophila* and mosquitos, germline sources of Cas9 are available ((Kondo and Ueda 2013; Gratz *et al*. 2014; Port *et al*. 2014; Li *et al*. 2017)). A key limiting factor in generating an endogenous source of Cas9, and in particular a source limited to the germline, is the identification of enhancers and promoters for tissue specific expression, a problem that is exacerbated in emerging animal models lacking extensive characterization of such regulatory elements. Despite the presence of homologs to elements validated in other insects, there remain few well-characterized and tested promoters for tissue-specific expression in *Tribolium* ((Lai *et al*. 2018; Khan *et al*. 2021)). Alternatively, a proven methodology has been to utilize a basal promoter to drive low ubiquitous expression (e.g., heat shock protein promoters). Indeed, on the available Cas9 plasmid ((Gilles *et al*. 2015)) for *Tribolium*, Cas9 is expressed from a core portion of the *Tribolium hsp68* promoter ((Schinko *et al*. 2010)) and does not require a heat pulse to induce editing ((Gilles *et al*. 2015; Rylee *et al*. 2018)).

Here, we report our creation of two *in vivo* endogenous sources of Cas9 facilitating general purpose genome modification in *Tribolium castaneum*. Utilizing the established expression of Cas9 from the *Tribolium hsp68* promoter ((Gilles *et al*. 2015)), we engineered a piggyBac transposable element carrying the previously described Cas9 cassette with a retinal GFP marker for screening (pB-hs-Cas9-hs). In the second piggyBac element, Cas9 is fused to EGFP via the viral T2A peptide, enabling spatiotemporal monitoring of expression, and the *hsp68* 3’UTR is exchanged for the *Tribolium nanos* 3’ UTR, along with a retinal *vermilion* marker for screening (pB-hs-Cas9-GFP-nanos). For each cassette, we isolated a transgenic line that was homozygous viable. Subsequent injection of gRNAs resulted in specific phenotypic and molecular modifications of the host genome, demonstrating the functionality of endogenous Cas9. These transgenic lines will increase the accessibility and feasibility of genome modification in *Tribolium*.

## Materials and Methods

### Tribolium husbandry and strains

All animals were raised at 28°C on a standard flour yeast mix. The following strains were utilized: *vermilion*^*white*^ (*v*^*w*^), ((Lorenzen *et al*. 2002)), m26, a *v*^*w*^ line with a X-linked insertion of the piggyBac transposase marked with 3XP3-DsRed ((Lorenzen *et al*. 2007)), piggyBac *3XP3-GFP; hsp68-nls-Cas9-nls-hsp3’UTR* (line #5556 (this study)), and *piggyBac 3XP3-v*^*+*^; *hsp68-nls-Cas9-nls-T2A-EGFP-nanos3’UTR* (line #3231(this study)). Injections were performed at 25°C and embryos were then returned to 28°C for development and hatching.

### Vectors

All vectors generated in this study are available through the Drosophila Genomics Resource Center at Indiana University.

#### pB-hs-Cas9-hs: *piggyBac 3XP3-GFP; hsp68-nls-Cas9-nls-hsp3’UTR*

The hsp68-nls-Cas9-nls-hsp3’UTR cassette was excised from p(bhsp68-Cas9) ((Gilles *et al*. 2015) Addgene (#65959)) using flanking AscI sites and ligated into AscI-linearized pBac-3XP3-GFP ((Horn and Wimmer 2000)).

#### pB-hs-Cas9-GFP-nanos: *piggyBac 3XP3-v*^*+*^; *hsp68-nls-Cas9-nls-T2A-EGFP-nanos3’UTR*

T2A-EGFP was amplified from pX458 (a gift from Feng Zhang, Addgene #48138), nanos3’UTR was amplified from *v*^*w*^ genomic DNA, and the fragments were assembled into p(bhsp68-Cas9) using NEBuilder HiFi assembly (NEB, Ipswich, MA). The hsp68-nls-Cas9-nls-T2A-EGFP-nanos3’UTR cassette was excised using flanking AscI sites and ligated into AscI-linearized pBac-3XP3-*v*^*+*^ ((Siebert *et al*. 2008)).

#### pU6b-BsaI-gRNA GFP1 and GFP2

The 20-mer protospacer sequences 5’-GGATCCACCGGTCGCCACCA-3’ and 5’-AAGGGCGAGGAGCTGTTCAC-3’ were cloned into BsaI-digested the pU6b-BsaI-gRNA ((Gilles *et al*. 2015), Addgene (#65956)) by NEBuilder HiFi-mediated ssDNA oligo bridging as described (https://www.neb.com/-/media/nebus/files/application-notes/bridging_dsdna_with_ssdna_oligo_and_nebuilder_hifi_dna_assembly_to_create_sgrna-cas-9_expression_vector_an.pdf?rev=6006359819ee4b06a269606b47110791) using an ssDNA oligo consisting of the protospacer flanked by 25bp regions of homology.

#### pU6b-BsaI-gRNA Tc vermilion

The 20-mer protospacer sequence 5’-GACCAACTGAGCGAAGAATG-3’ was cloned into BsaI-digested pU6b-BsaI-gRNA ((Gilles *et al*. 2015), Addgene (#65956)) by NEBuilder HiFi-mediated ssDNA oligo bridging as above using an ssDNA oligo consisting of the protospacer flanked by 25bp regions of homology.

### Tribolium transgenesis

pB-hs-Cas9-hs and pB-hs-Cas9-GFP-nanos were each resuspended in 1x injection buffer (0.5 mM KCl; 0.01 mM NaPO4 buffer pH 7.5) at a concentration of 1 µg/µL and injected into *m26* embryos. Individual surviving g^0^ adults were crossed to *v*^*w*^ and progeny were screened for expression of GFP in the retina or pigmented eyes, depending upon the injected construct. In total, three independent lines were isolated from pB-hs-Cas9-hs injections, but only one was capable of achieving homozygosity (Line # 5556). Two independent lines were isolated from pB-hs-Cas9-GFP-nanos and only one was capable of achieving homozygosity (Line # 3231).

### Tribolium CRISPR injection and detection

A mixture consisting of both GFP sgRNA plasmids (500 ng/µL of each) in 1x injection buffer (0.5 mM KCl; 0.01 mM NaPO_4_ buffer pH 7.5) was injected into embryos of line #5556. Individual surviving g^0^ adults were first screened to confirm the presence of 3XP3-GFP expression in the retina before individually crossing to *v*^*w*^. Individual g^0^ males were mated to 2-3 *v*^*w*^ females and individual g^0^ females were mated to 2-3 *v*^*w*^ males. Progeny were then subsequently screened for the loss of GFP expression in the retina. The progeny from individual crosses that lost GFP expression were saved and subjected to PCR and sequence confirmation for CRISPR/Cas9 editing.

The sgRNA plasmid for *vermilion* (500 ng/µL) was resuspended in 1x injection buffer (0.5 mM KCl; 0.01 mM, NaPO4 buffer pH7.5) was injected into embryos of line #3231. Individual g^0^ males were mated to 2-3 *v*^*w*^ females and individual g^0^ females were mated to 2-3 *v*^*w*^ males. Progeny were then subsequently screened for the loss of pigment in the retina. The progeny from individual crosses that lost pigment were saved and subjected to PCR and sequence confirmation for CRISPR/Cas9 editing.

### PCR and Sequence Confirmation of CRISPR/Cas9 editing

Genomic DNA was isolated from individual *Tribolium* by crushing each individual in 50µl of extraction buffer (100mM Tris-HCl, 50mM EDTA, 1% SDS) with a pestle in an Eppendorf tube. The mixture was subjected to a five-minute incubation at 95°C, then chilled on ice. The mixture was then digested with Proteinase K (50 µg/mL) for 1 hour at 55°C, followed by heat inactivation at 95°C for 5 minutes. 200µl of 0.1X TE buffer was added to dilute the sample. Finally, 100µl of the gDNA solution was purified using the Zymo Genomic DNA Clean and Concentrator –10 kit (Zymo Research #ZD4010) following the manufacturer’s instructions.

Amplicons spanning the gRNA target sites were amplified from 1 µL of purified gDNA using HotStar PCR Master Mix (Qiagen). Half of each reaction was run on a 1.5% agarose gel, and the other half of the reaction was purified using the Qiaquick Gel Extraction Kit (Qiagen). The purified fragments were submitted to Eurofins Genomics for Sanger sequencing, and the sequences were analyzed using Sequencher (Gene Codes Corp.). Alignments were made using SnapGene. Additionally, a fragment of Cas9 was amplified to confirm the presence of the 3XP3-GFP; hsp68-nls-Cas9-nls-hsp3’UTR insertion.

The following PCR primers were used to amplify the GFP and Cas9 DNA fragments: GFP – 5’-GCAAATAAACAAGCGCAGCTG-3’ and 5’-GTAGGTGGCATCGCCCTC-3’ and should amplify a 211 bp fragment. Cas9 - 5’-GGGATAAGCAATCTGGCAAA-3’ and 5’-CACGTGCCATTTCAATAACG-3’ and should amplify a 285 bp fragment. The GFP PCR products were sequenced with the following primer: 5’-CAAGCGCAGCTGAACAAGC-3’.

The following primers were used to amplify the *vermilion* DNA flanking the targeted gRNA site: 5’-ATGAGTTGCCCACTGAGACCCTCGTAAGTA-3’ and 5’-TGTTTGAACCATAACTCATACGCTGGAATTAC-3’ and should amplify a 297 bp fragment. The vermillion PCR products were sequenced with the following primers: 5’-GAGACCCTCGTAAGTATGATTTAA-3’ and 5’-CGCTGGAATTACTTGTTATAG-3’.

## Results and Discussion

### Generation of transgenic Cas9 T. castaneum lines

To streamline CRISPR genome editing in *T. castaneum*, we sought to generate lines in which the Cas9 coding sequence is stably integrated into the genome. To this end, we excised Cas9 under the control of the basal *hsp68* promoter from the original *T. castaneum* Cas9 vector bhsp68-Cas9 ((Gilles *et al*. 2015)) and inserted it into a piggyBac vector containing a 3XP3-GFP marker for random genomic integration ((Horn and Wimmer 2000)). The uninduced basal *hsp68* promoter and associated 3’ UTR have previously been demonstrated to provide sufficient Cas9 expression for genome editing in *Tribolium* ((Gilles *et al*. 2015; Rylee *et al*. 2018)). Subsequently, we sought to spatially restrict the activity of Cas9 to the posterior region of the embryo by exchanging the hsp68 3’UTR with the 3’UTR of *Tribolium nanos*. The 3’UTR of *Drosophila nanos* is critical for both localization and translation at the embryonic posterior pole ((Gavis and Lehmann 1992; Gavis and Lehmann 1994)). Nonetheless, whereas the function of *Tribolium nanos* appears to be conserved ((Schmitt-Engel *et al*. 2012)), whether similar functions of the 3’ UTR are conserved for *Tribolium nanos* remain unknown. As a means to monitor Cas9 expression from this construct, we also fused EGFP to Cas9 via the T2A peptide. The resulting construct was placed into a piggyBac vector containing a 3XP3-*v*^*+*^ as a marker for transgenesis ((Siebert *et al*. 2008)).

Both constructs, pB-hs-Cas9-hs and pB-hs-Cas9-GFP-nanos, were injected into *m26* embryos ((Lorenzen *et al*. 2007)) and surviving individual males and females were outcrossed to *v*^*w*^ beetles. To identify pB-hs-Cas9-hs insertions, progeny were screened for GFP expression in the retina (**Figure 1A**) and to identify pB-hs-Cas9-GFP-nanos insertions, progeny were screened for the restoration of pigment in the eye (**Figure 1B**). We isolated three independent lines for pB-hs-Cas9-hs and two for pB-hs-Cas9-GFP-nanos; however, only one line of each reached homozygosity, 5556 and 3231, respectively.

**Figure 1:**
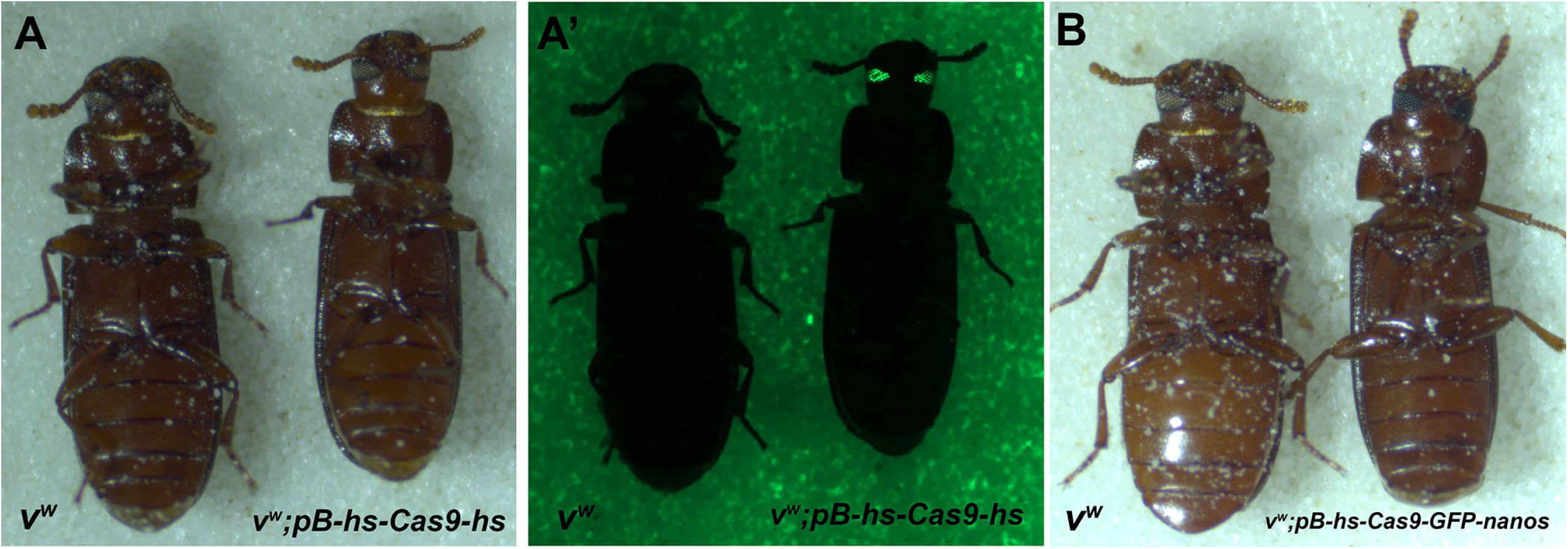
*Tribolium castaneum* genetic eye markers of Cas9 transgenic lines. **A, A’**. Comparison of *v*^*w*^ to pB-hs-Cas9-hs (*piggyBac 3XP3-GFP; hsp68-nls-Cas9-nls-hsp3’UTR*). Note both contain the loss of eye pigment due to the presence of *v*^*w*^, the loss of *vermillion*, but pB-hs-Cas9-hs is distinguished by GFP (3XP3-GFP) expression in the retina. B. Comparison of *v*^*w*^ to pB-hs-Cas9-GFP-nanos (*piggyBac 3XP3-v*^*+*^; *hsp68-nls-Cas9-nls-T2A-EGFP-nanos3’UTR*). Note the pigmented eyes in pB-hs-Cas9-GFP-nanos due to the transgene rescue (3XP3-*v*^*+*^) of *vermillion*.

### Transgenic Cas9 is functional

To test the inserted pB-hs-Cas9-hs for functionality, we made use of the 3XP3-GFP region of the integrated piggyBac transposon. We injected plasmids expressing two different sgRNAs flanking the start codon of EGFP, theoretically leading to indels that would disable GFP translation initiation (**Figure 2A**). 110 g^0^ embryos survived to adults and were individually outcrossed to *v*^*w*^ beetles. The progeny of each cross was screened for loss of GFP expression in the eye. Genomic DNA was collected from beetles lacking GFP expression, and a region spanning the predicted mutation site, as well as a short span in the Cas9 transgene, were PCR amplified. Eleven crosses yielded progeny with no GFP expressed in the eye. Of these, amplification of both GFP and Cas9 failed using genomic DNA recovered from progeny of nine crosses, suggesting a lack of transgene integration. In addition, these nine crosses generated an approximate ratio of 1:1 with respect to the absence or presence of GFP. We attribute the lack of GFP to a low level of heterozygosity of the pB-hs-Cas9-hs in our 5556 stock rather than variable Cas9 activity. We recovered amplification products for both GFP and Cas9 in the two remaining lines that yielded GFP-negative progeny, F14 and F102. Sequencing of the GFP PCR product (**Figure 2B and 3B, Supplemental Figure 1A**) revealed F14 progeny contained a complex indel within the gRNA-1 site, and a 5 bp deletion within the gRNA-2 site (**Figure 1C**), suggesting both sgRNAs were capable of mediating Cas9 directed editing. In contrast, the F102 progeny contained a 96 bp insertion within the gRNA-1 site (**Figure 3C, Supplemental Figure 1B**); the sequence of the 96 bp is identical to a region of pU6b-BsaI-gRNA vector.

**Figure 2:**
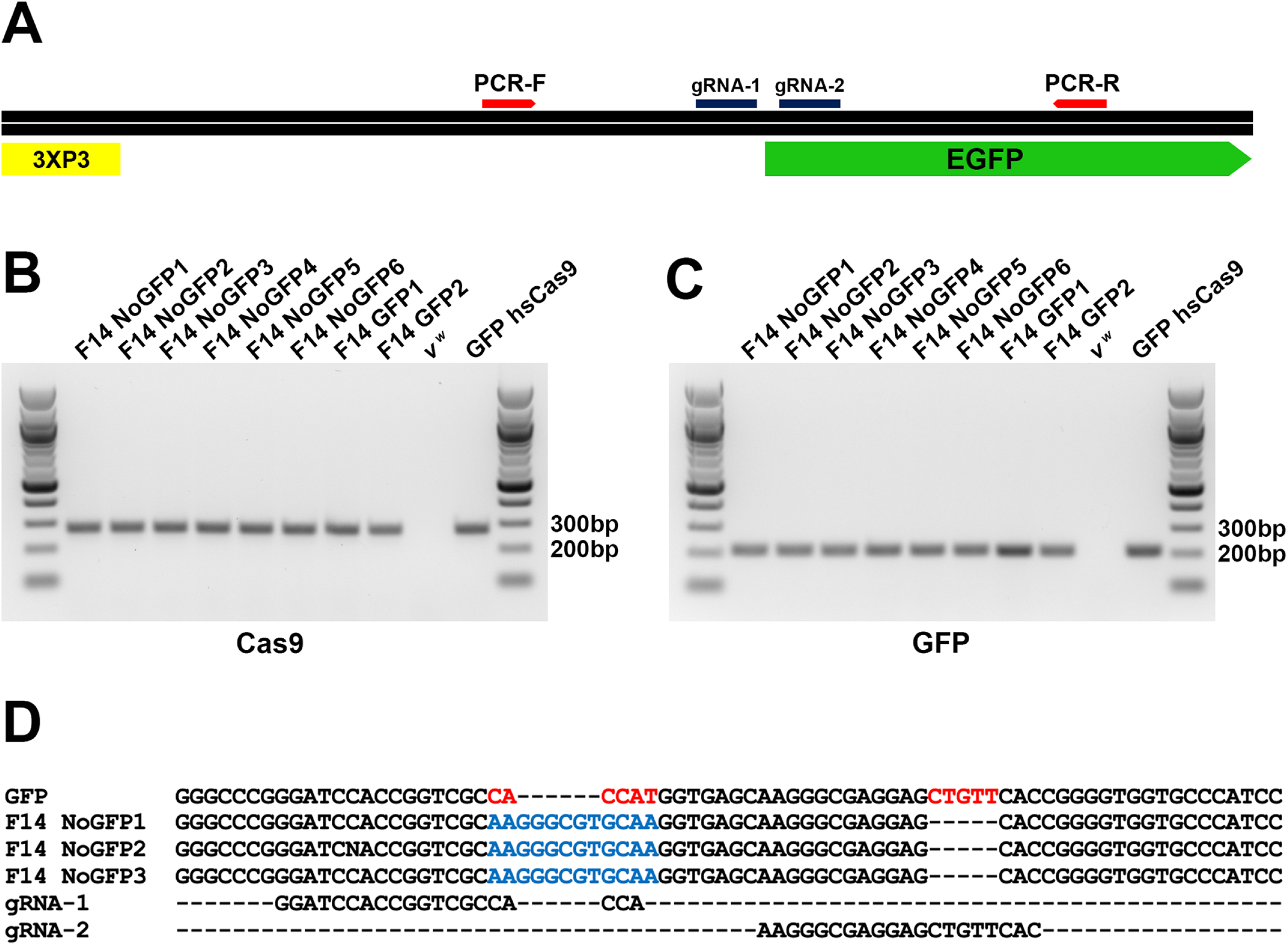
Characterization of CRISPR/Cas9 induced mutation in progeny of female-14. **A**. Schematic showing the locations of gRNA targets, primers for PCR amplification, EGFP, and the 3XP3 promoter within the transgenic insertion. **B-C**. Gel images of PCR amplification products of Cas9 (**B**) or GFP (**C**) from genomic DNA extracted from individual progeny with or without GFP expression visible in the retina, *v*^*w*^, and parental 5556 beetle containing the pB-hs-Cas9-hs construct. **D**. Alignment of sequences from GFP amplification products from progeny lacking GFP expression as compared to the parental pB-hs-Cas9-hs transgenic beetle; red bases indicate bases deleted and blue bases represent insertions as compared to the wild type.

**Figure 3:**
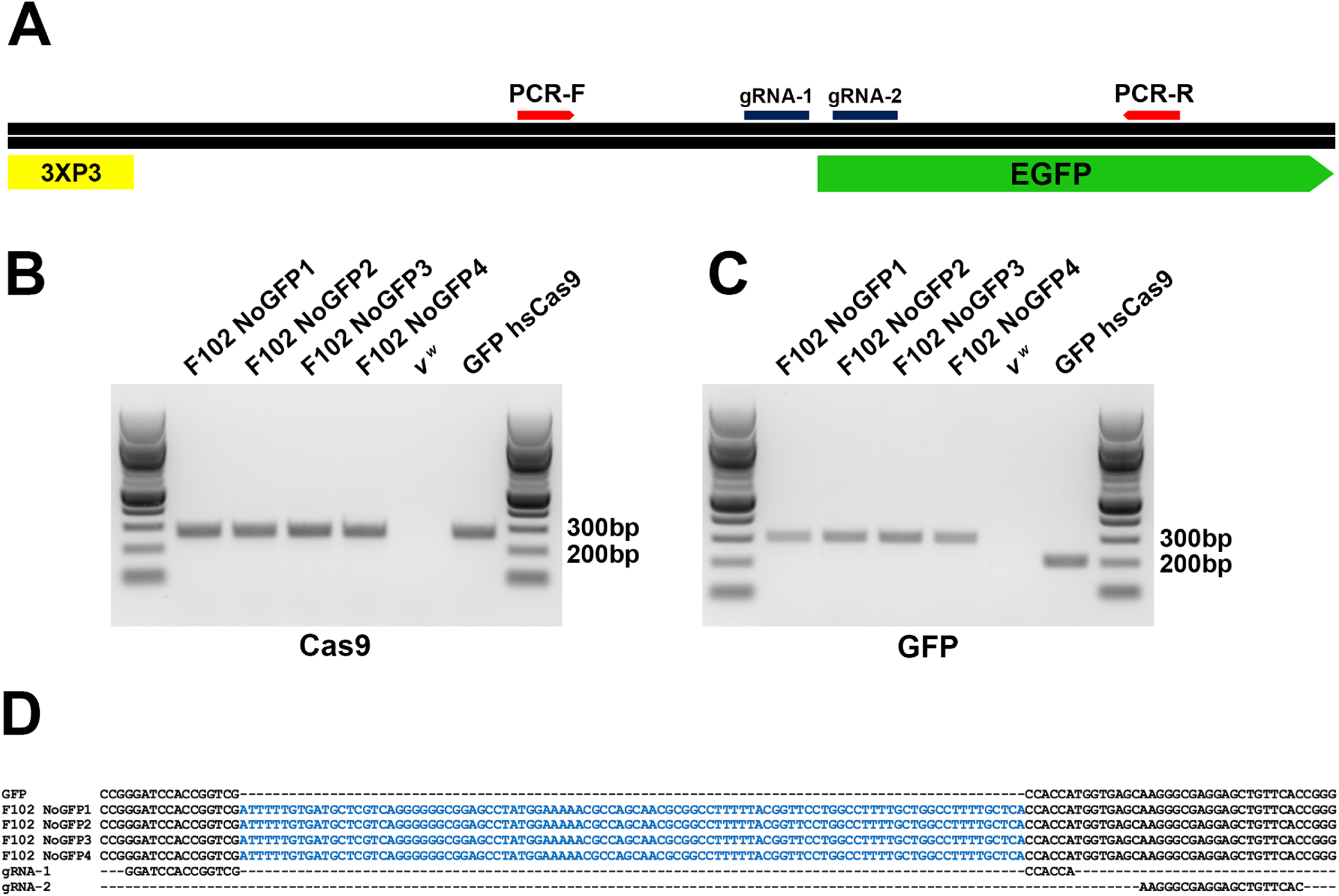
Characterization of CRISPR/Cas9 induced mutation in progeny of female-102. **A**. Schematic showing the locations of gRNA targets, primers for PCR amplification, GFP, and the 3XP3 promoter within the transgenic insertion. **B-C**. Gel images of PCR amplification products of Cas9 (**B**) or GFP (**C**) from gDNA extracted from individual progeny with or without GFP expression visible in the eyes, *v*^*w*^, and parental 5556 beetle containing the pB-hs-Cas9-hs construct. **D**. Alignment of sequences from GFP amplification products from progeny lacking GFP expression as compared to the parental pB-hs-Cas9-hs transgenic beetle; blue bases represent insertions as compared to the wild type.

Similar to the first construct, for pB-hs-Cas9-GFP-nanos, we targeted the wild-type *vermilion* gene within the *piggyBac 3XP3-v*^*+*^ inserted cassette for editing. As previously reported, the *v*^*w*^ strain contains a partial deletion of the *vermilion* gene wherein only the 3’ end of the gene is present ((Adrianos *et al*. 2018)). Hence, our sgRNA targets the second exon of the *vermilion* transgene present in the piggyBac transposable element (**Figure 4A**). The sgRNA was injected into a mixed stock of heterozygous and homozygous beetles of line #3231. We recovered 87 g^0^ injected adults and upon subsequent mating to *v*^*w*^ demonstrated 40 of the surviving g^0^ adults were heterozygous for the piggyBac element based upon the expected Mendelian ratio of 1:1 of pigmented to non-pigmented eyes in the progeny. Therefore, the progeny of these 40 crosses were not examined for any potential Cas9 editing. 4 g^0^ crosses resulted in no progeny and out of the 43 remaining crosses we recovered one g^0^ male cross that contained 2 progeny that lacked eye pigmentation among many pigmented siblings. PCR amplification (**Figure 4B**) and sequencing of these two progeny revealed and confirmed a single base pair deletion localized to the sgRNA target site (**Figure 4C and Supplemental Figure 1C)**. The deletion results in a premature stop codon in the second exon of the transgene thus eliminating a functional protein (**Figure 4C**). These results indicate that both construct/transgenic lines are capable of inducing Cas9 mediated genome editing. For each construct the mutagenesis rates that we observed were relatively low. This may be due to the basal expression levels provided by the *hsp68* core promoter used in both of our expression cassettes. It was reported that the *hsp68* core promoter gives only basal levels of expression, while expression is significantly higher when an approximately 450 bp region upstream of the core promoter was included ((Schinko *et al*. 2010)). In addition, our results do not permit a comparison between mutagenesis rates of constructs, given our experiments were designed only for testing functionality and different sgRNAs and target genes were utilized in each case.

**Figure 4:**
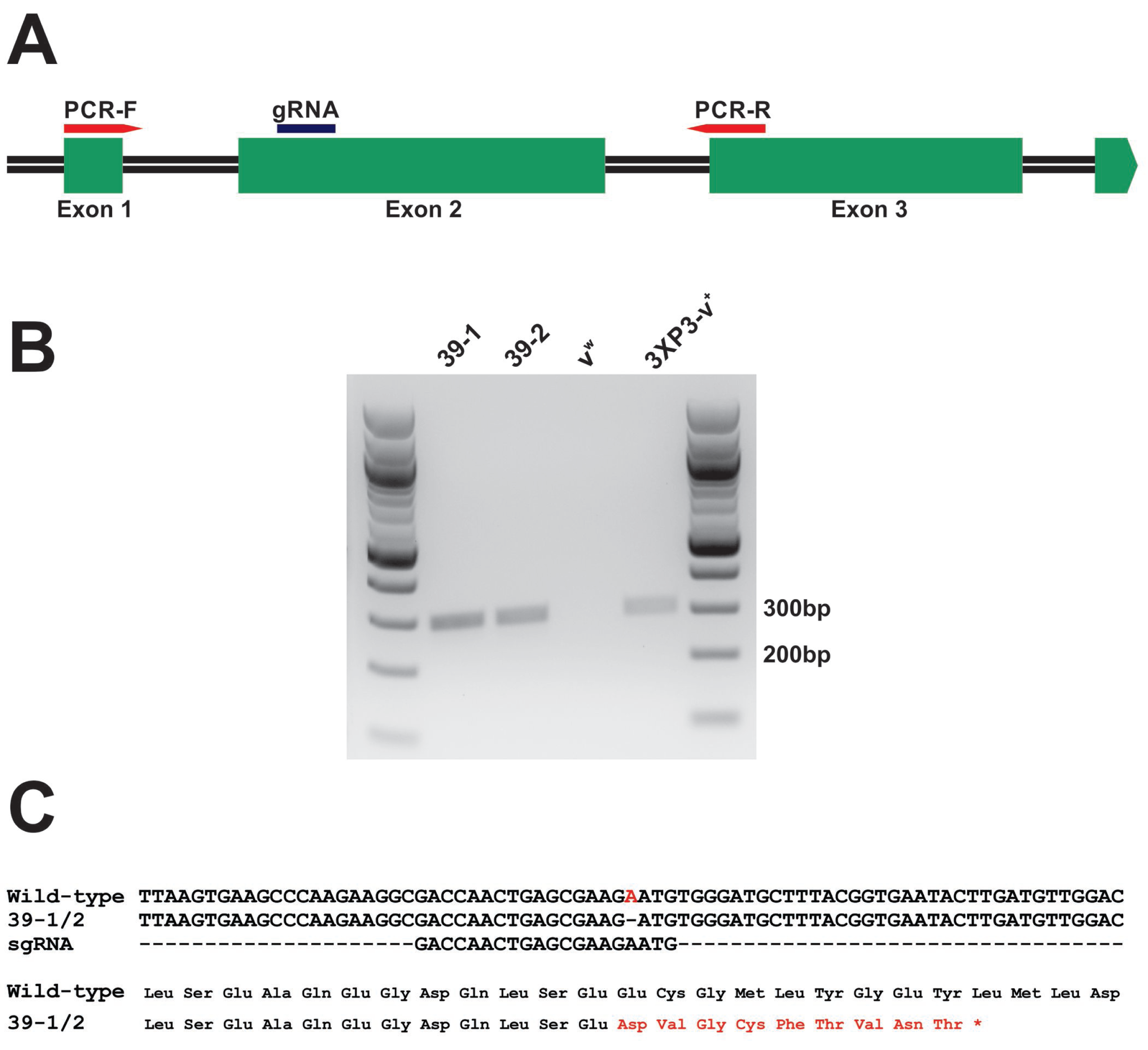
Characterization of CRISPR/Cas9 induced mutation in *vermillion* transgene. **A)** Schematic showing the locations of the gRNA target and PCR primers within the first three exons of the *vermillion* transgene. **B**) Gel image of the amplification products. **C**) Alignment of sequences and protein translation from a pB-hs-Cas9-GFP-nanos, and from animals 39-1 and 39-2. Red bases indicate bases that are deleted in the mutated beetles.

However, increasing ubiquitous expression of Cas9 could result in toxicity. With respect to potential toxicity of expressing low levels of Cas9, we can only comment on the observed different rates of g^0^ crosses resulting in no progeny. In line 5556, pB-hs-Cas9-hs, the construct significantly affected male fecundity; 17 of the 54 g^0^ males did not produce progeny whereas 5 of the 47 g^0^ females did not produce progeny (**Table 1**). In the case of line 3231, pB-hs-Cas9-GFP-nanos, we observed the opposite. Four of the 20 g^0^ female crosses did not produce progeny, whereas all 27 g^0^ male crosses produced progeny (**Table 1**), a significant reduction in female fecundity. Male reproduction varied significantly between the two constructs (p < 0.004) but female reproduction did not (p > 0.25). Although these data are consistent with the suggestion that the female-specific nanos 3’UTR is less harmful to male reproduction, our current results cannot distinguish whether the differences observed are due to Cas9 activity, insertion location of transgenes, or differences in constructs. For example, due to low expression from the *hsp68* core promoter we were not capable of utilizing GFP expression as a surrogate for Cas9 activity in the pB-hs-Cas9-GFP-nanos transgenic line 3231.

**Table 1:** Fecundity comparisons of males and females with respect to transgenic Cas9 construct.

*piggyBac 3XP3-GFP; hsp68-nls-Cas9-nls-hsp3'UTR*
| | **Males** | **Females** | $\Sigma$ |
| --- | --- | --- | --- |
| **Yielded Progeny** | 37 | 42 | 79 |
| **No Progeny** | 17 | 5 | 22 |
| $\Sigma$ | 54 | 47 | 101 |
|  |  |  | **p = 0.0152\*** |

\*Fisher's Exact Test, significant difference between males and females.

*piggyBac 3XP3- $v^{w+}$ ; hsp68-nls-Cas9-nls-T2A-EGFP-nanos3'UTR*
| | **Males** | **Females** | $\Sigma$ |
| --- | --- | --- | --- |
| **Yielded Progeny** | 27 | 20 | 47 |
| **No Progeny** | 0 | 4 | 4 |
| $\Sigma$ | 27 | 24 | 51 |
|  |  |  | **p = 0.0272\*** |

\*Fisher's Exact Test, significant difference between females and males.

Altogether, we have generated the first *Tribolium* lines expressing Cas9 endogenously, adding to the ever-growing set of genome editing tools available to the community. Our work demonstrates the feasibility of these resources and also sets a baseline against which to compare other promoter-Cas9 combinations. As observed in other insect species, further work is needed to identify and characterize usable regulatory regions to drive robust germline Cas9 expression ((Kondo and Ueda 2013; Gratz *et al*. 2014; Li *et al*. 2017)). Nonetheless, these transgenic Cas9 lines will increase the accessibility of CRISPR/Cas9 editing in *Tribolium* for the research community.

## Data availability

All data necessary for confirming the conclusions in this paper are included in this article and in supplemental figures and tables.

## Acknowledgements

This work was supported by NIFA/USDA 2019-33522-30064 (M.J.W, A.C.Z., G.E.Z.) and NSF grant IOS-1928781 (A.C.Z.).

## Figure Legends

**Supplementary Figure 1: Chromatography readouts of CRISPR/Cas9 editing**.

**A)** Chromatograms from sequencing PCR products amplified from gDNA extracted from pB-hs-Cas9-hs control beetles, and the progeny from female 14 lacking GFP expression. **B)** Chromatograms from sequencing PCR products amplified from gDNA extracted from pB-hs-Cas9-hs control beetles, and the progeny from female 109 lacking GFP expression. The site of the insertion is shown **C)** Chromatograms from sequencing PCR products amplified from gDNA extracted from pB-hs-Cas9-GFP-nanos control beetles, and the progeny from 39-1 lacking eye pigment. Red boxes indicate deletions, and blue boxes indicate insertions.

